# The role of stimulus periodicity on spinal cord stimulation-induced artificial sensations in rodents

**DOI:** 10.1101/2022.12.26.521912

**Authors:** Jacob C. Slack, Sidnee L. Zeiser, Amol P. Yadav

**Author notes:** These authors contributed equally (Jacob C. Slack, Sidnee L. Zeiser). **Corresponding Author: Amol P. Yadav, Ph.D.**, Assistant Professor of Neurological Surgery, Stark Neurosciences Research Institute, Indiana University School of Medicine, Indianapolis, IN-46202, USA.

## Abstract

Sensory feedback is critical for effectively controlling brain-machine interfaces (BMIs) and neuroprosthetic devices. Spinal cord stimulation (SCS) is proposed as a technique to induce artificial sensory perceptions in rodents, monkeys, and humans. However, to realize the full potential of SCS as a sensory neuroprosthetic technology, a better understanding of the effect of SCS pulse train parameter changes on sensory detection and discrimination thresholds is necessary. Here we investigated whether stimulation periodicity impacts rats’ ability to detect and discriminate SCS-induced perceptions at different frequencies. By varying the coefficient of variation (CV) of interstimulus pulse interval, we showed that at lower frequencies, rats could detect highly aperiodic SCS pulse trains at lower amplitudes (i.e., decreased detection thresholds). Furthermore, rats learned to discriminate stimuli with subtle differences in periodicity, and the just-noticeable differences (JNDs) from a highly aperiodic stimulus were smaller than those from a periodic stimulus. These results demonstrate that the temporal structure of an SCS pulse train is an integral parameter for modulating sensory feedback in neuroprosthetic applications.

## Introduction

Neuroprosthetic devices and BMIs have successfully demonstrated restoration of motor function in individuals suffering from impairments caused by spinal cord injuries (SCIs) and limb amputations (1–3). However, methods that can restore tactile and proprioceptive functions via a non-visual pathway such as intracortical stimulation of the somatosensory cortex, deep brain stimulation, and stimulation of peripheral nerves are still under investigation (4–7). An alternative method, spinal cord stimulation (SCS), was recently shown to successfully generate artificial sensations in rats, monkeys, and amputee human subjects with limb amputation (8–10). Yet, exact spinal stimulation parameters that can reproducibly induce naturalistic sensory perceptions are still to be determined.

Traditional stimulation pulse trains are periodic in nature; however, the naturalistic pattern of neural activity representing tactile and proprioceptive signals is usually aperiodic (11). It is therefore argued that to mimic naturalistic sensations, it might be necessary to deliver pulse trains with a degree of aperiodicity or randomness (11, 12). Previously, it was shown that SCS-induced sensory detection thresholds decreased with increasing frequency, pulse width, and duration of stimulation (9). Moreover, although rats learned to discriminate SCS frequencies and frequency discrimination obeyed Weber’s law, the stimulation train used was always periodic in nature. Thus, it needs to be investigated whether rats can learn to detect and discriminate aperiodic SCS pulse trains and whether varying periodicity changes detection and discrimination thresholds. This will allow the determination of upper and lower limits for periodicity as novel temporal stimulation patterns are explored. In this study, we used a two-alternative forced choice (2AFC) task to determine sensory detection thresholds at different levels of aperiodicity determined by the coefficient of variation (CV) of the interstimulus pulse interval. We also determined the just-noticeable difference (JND) in aperiodicity that rodents can successfully discriminate at multiple stimulation frequencies.

## Methods

All animal procedures were approved by the Indiana University Institutional Animal Care and Use Committee (IACUC) and were performed in accordance with National Institute of Health Guide for the Care and Use of Laboratory Animals. Six Long Evans rats (250-400g) participated in the experiments (Supplementary Table 1).

### Pre-training and spinal implant surgery

Upon animal delivery, baseline weights of all rats were taken, and rats were acclimated to human handling for 1-2 days. After acclimation, rats were moderately water deprived and were placed in the operant training chamber for approximately one week to become comfortable with the experimental environment. The chamber contained two water dispensing reward ports on each side that were enclosed by doors and placed behind infrared (IR) beam sensors (Fig 1a). A water reward was delivered when the rat completed a nose poke into the port, which interrupted the IR beam. A light in the chamber would turn on to signify the start of each trial. Initially, the rats were rewarded with water after making a nose poke and licking at either spout. Following that, rats were trained to make nose pokes in each spout on alternate trials. The initial learning period took between 7-23 days, i.e. a consistent performance (>90%) on alternating reward port trials.

**Figure 1.**
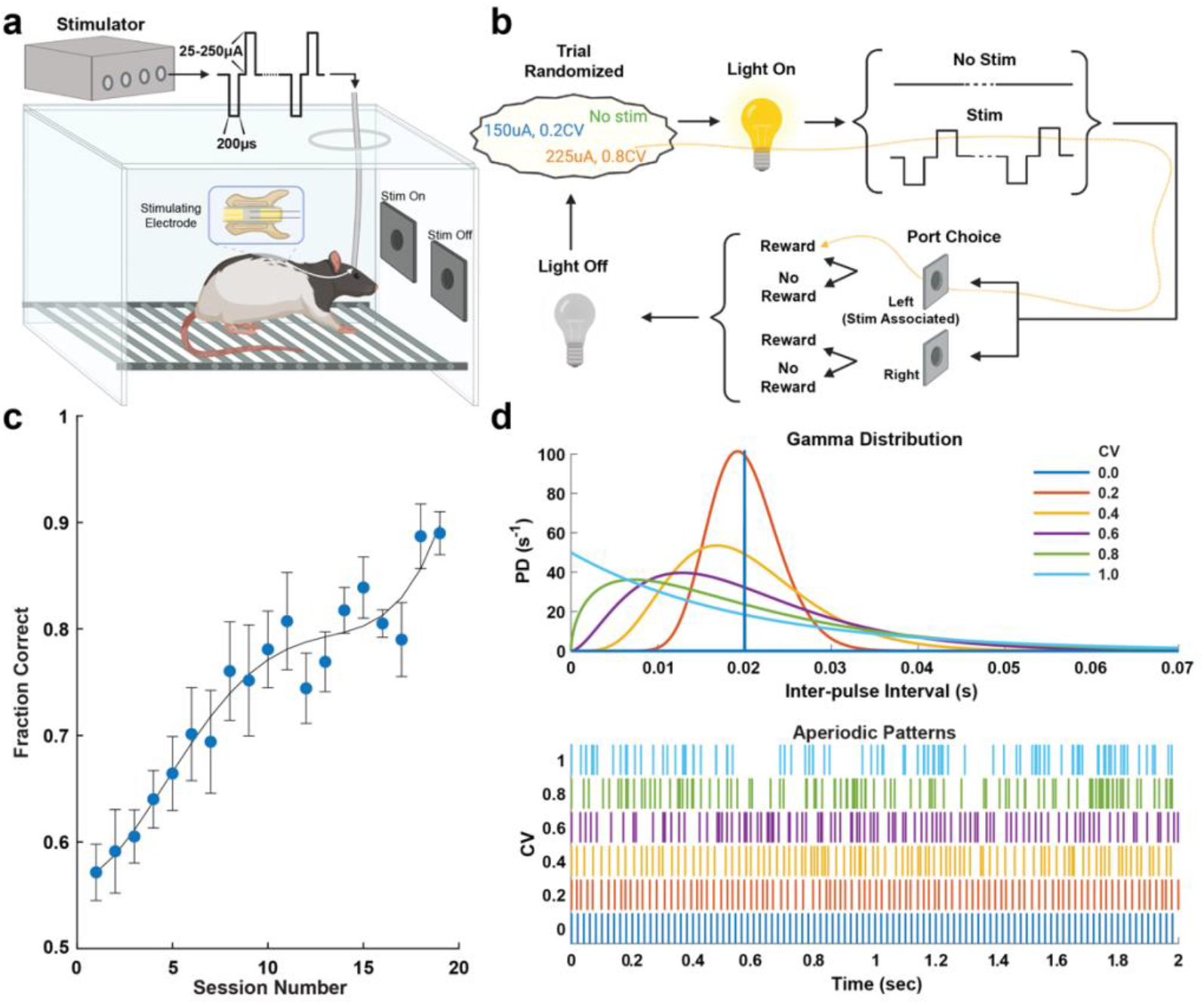
Behavioral task design and aperiodic SCS pulse-train detection learning. a) Behavioral chamber with two water reward ports covered by doors for conducting detection and discrimination experiments. A microstimulator delivered biphasic, bipolar, charge-balanced pulse trains across the two-lead epidural electrode in six freely moving rats. b) Task flow of a detection trial. A session normally consisted of 200-400 trials. ‘Stim’ and ‘No Stim’ trials were randomly selected followed by a light indicator followed by presentation of a sensory cue for two seconds. Rats received either SCS for ‘Stim’ trials or absence of SCS for ‘No Stim’ trials. Reward port doors would then open allowing the rat to select either left or right port. A water reward paired with a short auditory cue was delivered if a rat chose the left port during a ‘Stim’ trial or the right port during a ‘No Stim’ trial. If an incorrect port was chosen, no reward was delivered paired with a long auditory cue. c) Learning curve for initial detection training averaged across six rats. Fraction of correct trial decisions is plotted across sessions. Circles and error bars indicate mean±standard deviation. d) Aperiodic template patterns generated from a gamma distribution using the coefficient of variation (CV) of the interstimulus pulse interval to vary the level of periodicity (CV=0: highly periodic; CV=1: highly aperiodic).

Once rats successfully learned how to interact with the operant chamber, custom two-contact SCS platinum electrodes (1×0.5mm) spaced 0.25mm and 0.025mm thick with PFA coated stainless steel leads (A-M Systems, Sequim, WA, USA) were implanted epidurally underneath the T3-T5 vertebra using previously established procedures (9, 13–15). The leads were then routed to an Omnetics connector (Omnetics Connector Corporation, MN, USA) and fixed to the skull with skull-screws (W. W. Grainger Inc., IL, USA) and dental acrylic (Colten Holding, Switzerland). After recovery, electrodes were tested using a custom biphasic/bipolar microstimulator (16). Tests consisted of applying pulse-trains using stimulation parameters consistent with the experimental task (i.e, 50Hz pulse-train above sensory threshold but below motor threshold).

### Aperiodic SCS Pulse-Trains

SCS pulse-trains were bipolar, biphasic, and charge-balanced with 200μs phase duration and inter-phase interval of 50μs. To generate aperiodic SCS pulse-trains, inter-pulse interval values were pulled from a gamma distribution in MATLAB (MathWorks, MA, USA) using Eq. 1 (12).

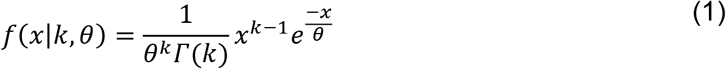

with probability density function *f*, inter-pulse interval *x* in milliseconds, the gamma function *Γ*, shape and parameter *k* and scale parameters θ. The degree of aperiodicity was modulated by changing the shape and scale parameters which are defined as follows:

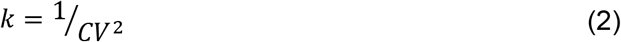

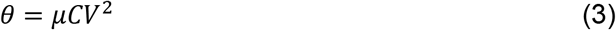

where CV is the coefficient of variation and *μ* the mean inter-pulse interval. Larger CV values shift the peak of the probability density function leftward producing increasingly aperiodic patterns (Fig 1d). For psychometric detection, six CV values were chosen ranging from 0 (periodic) to 1 incremented by 0.2. In discrimination experiments, ten CV values were used incremented by 0.1 from 0 to 1. In both cases, a template pattern was generated from Eq. 1 for each CV value at each frequency and used for all rats. The template pattern was limited to a two second window and contained equal pulse across CV values within a given frequency. As an example, the template patterns for 50Hz at 0.2 CV and 50Hz at 1 CV both contained 100 pulses within the two-second stim period.

### Sensory detection task and psychophysics

To learn the stimulation detection task, rats were trained on a 2AFC task. We used a behavioral chamber (Med Associates Inc., VT, USA) described above and previously (8, 9) but modified the electrical interface so that it could be custom-controlled by Arduino (Arduino LLC, Italy) and interfaced with MATLAB via a GUI (Fig 1a). Each trial consisted of a houselight turning on followed by stimulation present (‘Stim’ trial) or stimulation absent (‘No Stim’ trial) (Fig 1b). Trial selection was randomized programmatically. For initial detection training stimulation amplitude above sensory threshold but below motor threshold was determined before each session by the experimenter. Previously we used periodic pulse trains at 100 Hz for detection training, but here, the pulse-train (cathode leading with 200us pulse width, 50us interphase delay, 2s duration) was delivered at 0.8CV at 50 Hz. Once rats reached a correct detection rate of 80% (Fig 1c), CV (0–1) and amplitude (ranging from below sensory threshold to above sensory threshold) were randomized for ‘Stim’ trials at a given frequency (Fig 1d). When 20 trials for all CV and amplitude combinations were completed, a different frequency was selected, and the process repeated. Sensory detection thresholds were calculated as 75% of correct detections from sigmoid fits.

### Initial sensory discrimination task

The sensory discrimination task also utilized a 2AFC task setup and training chamber. Prior to the session, each individual rat’s sensory threshold amplitude was determined with a 2 second periodic (0 CV) pulse train. This amplitude was constant throughout the session unless the rat was performing poorly and needed an adjustment. The stimuli varied with either a high (1) or low (0) CV. Other parameters such as pulse width (200 us), amplitude (determined and set at the beginning of each session), frequency (20, 50, 100, or 200 Hz), duration (2 s) remained the same throughout the session. For the initial discrimination task, the high CV was associated with the left reward port while the low CV corresponded to the right. Once each rat consistently showed the ability to discriminate between 1 CV and 0 CV by achieving a correct response rate >80%, they were advanced to the next task to determine just-noticeable differences in CV.

### Just-Noticeable Difference and sensory discrimination

After learning the initial CV discrimination (0 vs. 1 CV), the left port was held at a constant 1 CV (standard CV) while the right port CV (comparison CV) was randomized from 0 to 0.9 in increments of 0.1. The same experiment was then repeated while keeping the right port at a constant 0 CV (standard CV) and randomizing the left port CV (comparison CV) from 0.1 to 1. The standard CV was not changed during a particular session. Trial type and comparison CV were randomized by a programmatic algorithm. Discrimination performance was calculated as the fraction of total correct trials for each standard and comparison pair presentation. The fractions were plotted against their corresponding standard/comparison CV difference. A sigmoidal curve was then fitted to the points. The Just-Noticeable Difference (JND) of CV was determined as the 75% correct percentage mark on the resulting curve. Once JND was determined for a particular frequency, the same procedures (initial discrimination training followed by CV randomization) were repeated at different frequency values to study additional trends.

### Statistical Analysis

Before performing statistical analysis, thresholds were normalized by the maximum threshold for each rat. Repeated Measures one-way ANOVA with post-test for trend and multiple comparisons was applied to determine significant differences between thresholds at various degrees of periodicity.

## Results

### Sensory Detection of Aperiodic SCS Pulse-Trains

During initial training, six rats learned to detect cathode-leading, biphasic, aperiodic pulse-trains (frequency: 50Hz, CV: 0.8, pulse-width: 200ms, inter-phase delay: 50μs, amplitude: 190.8±90.1μA, Fig. 1c) over the course of 11 ±5 sessions. To determine sensory detection thresholds at different CV values (0 to 1), amplitude and CV were randomized at each stimulation frequency (10-200 Hz), and psychometric curves were fit to the resultant detection performance values (Fig 2a). Averaged detection thresholds across all rats showed that detection thresholds were significantly different between 0 CV and 1 CV stimuli at 50 Hz (Fig 2c, third column, p < 0.01, repeated measures (RM) one-way ANOVA multiple comparisons). Detection thresholds were also found to have a decreasing linear trend for both 20Hz (Fig 2c, second column, p < 0.05, RM one-way ANOVA test for trend) and 50Hz (Fig 2c, third column, p < 0.05, RM one-way ANOVA test for trend) suggesting that rats detected SCS at a lower amplitude as aperiodicity of the stimulus train (CV) was increased.

**Figure 2.**
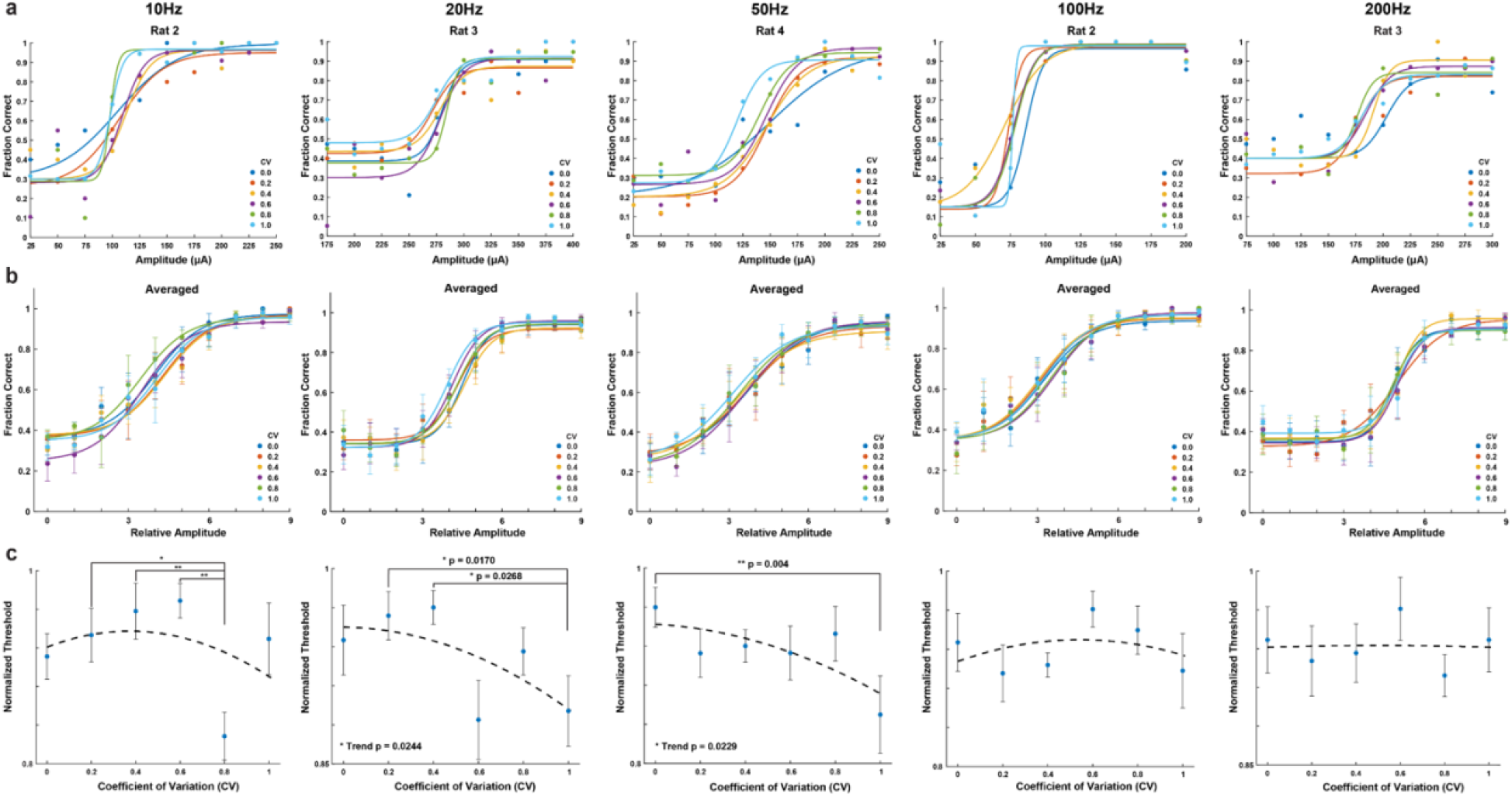
Psychometric analysis of SCS sensory detection for various degrees of aperiodicity at constant frequency. Columns indicate the frequency of SCS pulse-trains. a) Representative psychometric detection curves for individual rats at multiple CV values at constant frequency. b) Psychometric detection curves averaged across rats for each frequency (10, 20, 50, 100, 200 Hz; n=3, 6, 6, 4, 3). X-axis depicts amplitude relative to individual rats (amplitude ranges subtracted by minimum amplitude and divided by step size). Circles and error bars indicate mean ± standard error. Traces for ‘a’ and ‘b’ are sigmoid fits for individual CV values across amplitudes. c) Normalized detection thresholds across CV for constant frequency. For each rat, thresholds were determined from individual sigmoid fits as seen in ‘a’ as the 75% fraction of correctly detected stimuli and normalized by maximum amplitude. Circles and error bars indicate mean ± standard error. P-values were calculated from repeated measures one-way ANOVA post-test for trend and multiple comparisons.

Analyzing detection performance across multiple frequencies demonstrated that detection thresholds decreased with increasing frequency for all CV values (Fig. 3, p < 0.001, one-way ANOVA). This confirmed a previous result where detection thresholds decreased with increasing frequency (9), however, in that case, the stimulus pulse train was only delivered in a periodic form (corresponding to 0 CV here). Thus, our results establish that the relationship between detection thresholds and frequency is maintained independently of stimulus periodicity.

**Figure 3.**
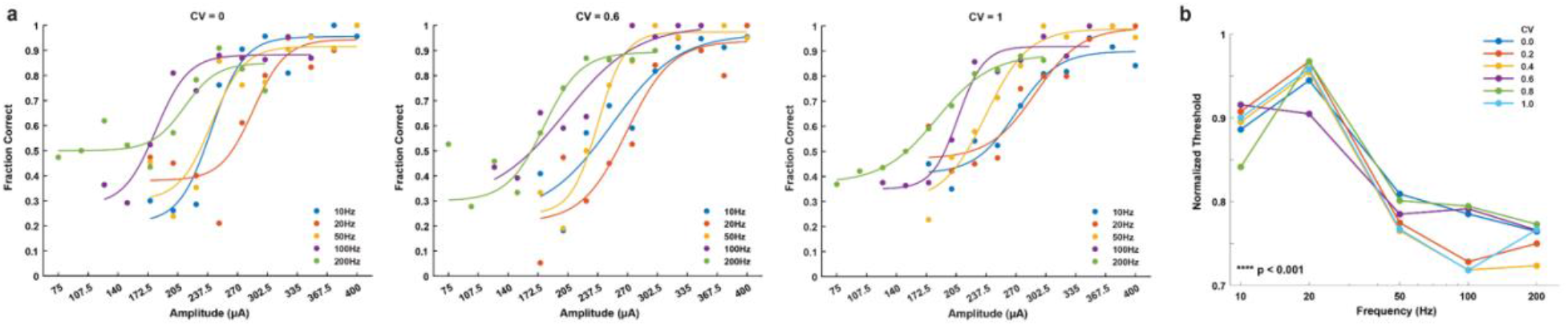
Psychometric analysis of sensory detection across frequency. a) Representative psychometric detection curves across frequency for rat 3. Traces are sigmoid fits for multiple frequencies at a given CV value. b) Normalized detection thresholds across frequency. Traces indicate normalized thresholds for a CV value at each frequency. P-value was calculated by one-way ANOVA.

### Sensory Discrimination of Aperiodic SCS Pulse-Trains

Following the detection behavioral task, four rats were trained to discriminate between SCS patterns that varied in CV while the amplitude, frequency, pulse-width, and duration were kept constant throughout a session. All four rats learned the initial task of discriminating between a CV of 1 and CV of 0; rat 1 and 4 successfully learned at 50 Hz while rat 2 and 3 successfully learned at 20 Hz (Supplementary Fig 1). The rats achieved a consistent correct percentage rate ≥ 80% between 12-34 days (Fig. 4).

**Figure 4.**
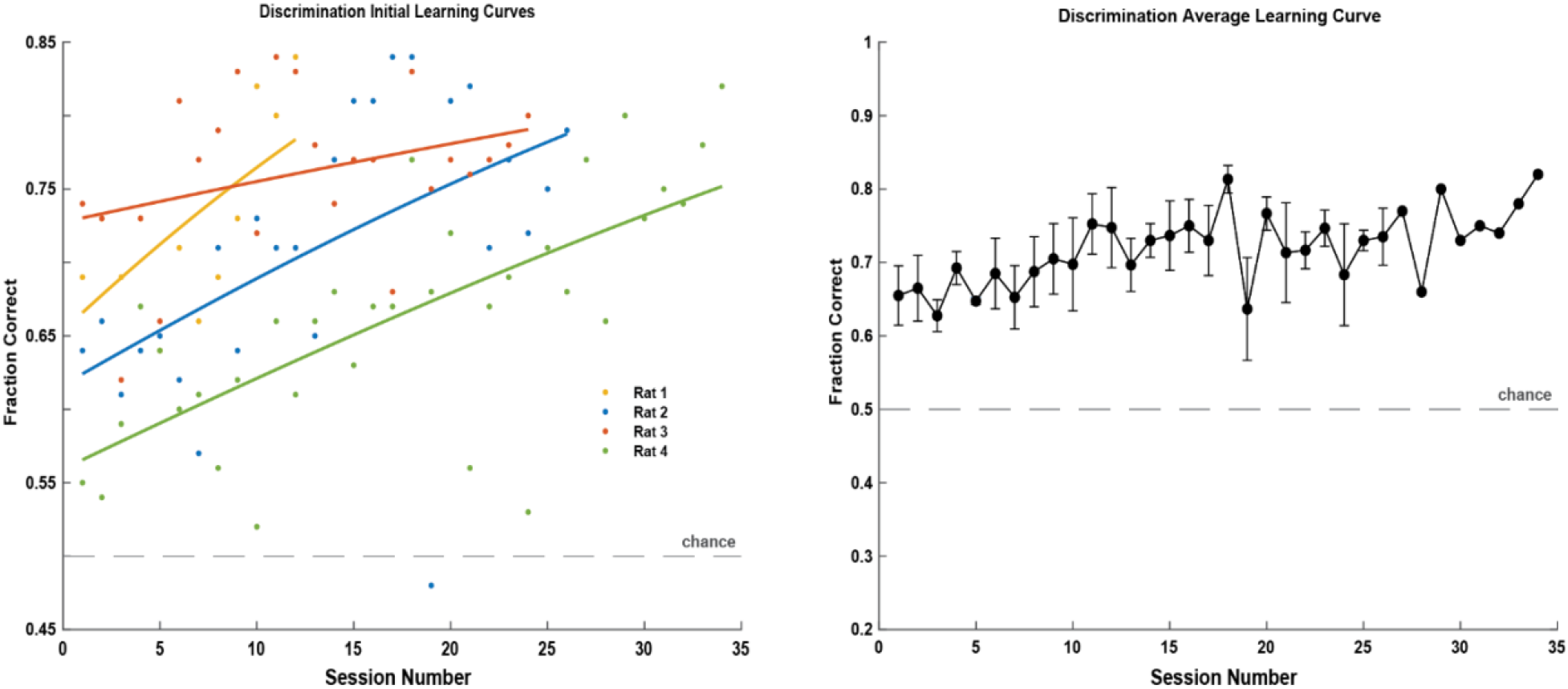
Learning curves for the initial discrimination task. a) Four rats learned to discriminate SCS stimuli with a CV of 0 versus a CV of 1 within 12-34 days. Rats 1 and 4 learned at 50 Hz, while rats 2 and 3 learned at 20 Hz. The learning curves show the trial-average fraction of correct trials as a function of training sessions. b) The average fraction of correct trials for the four rats. Circles and error bars indicate mean ± s.e.m.

Once rats learned to successfully discriminate between highly periodic (0 CV) and highly aperiodic (1 CV) stimuli, we determined JNDs by keeping standard CV of 1 while varying comparison CV from 0-0.9. Our results showed that rats were able to discriminate from a highly aperiodic stimulus at all tested frequencies (20, 50, 100, and 200 Hz), and the JNDs in CV for successful discrimination were 0.27 at 20 Hz, 0.5 at 50 Hz, 0.49 at 100 Hz, and 0.87 at 200 Hz across all rats (Fig 5a, 6a). To determine JNDs from a periodic stimulus we kept standard CV of 0 and varied the comparison CV from 0.1-1 for all frequencies (Fig 5b). JNDs in CV for successful discrimination were 0.85 at 20 Hz, 0.86 at 50 Hz, 0.73 at 100 Hz, and 0.69 at 200 Hz across all rats (Fig 6b).

**Figure 5.**
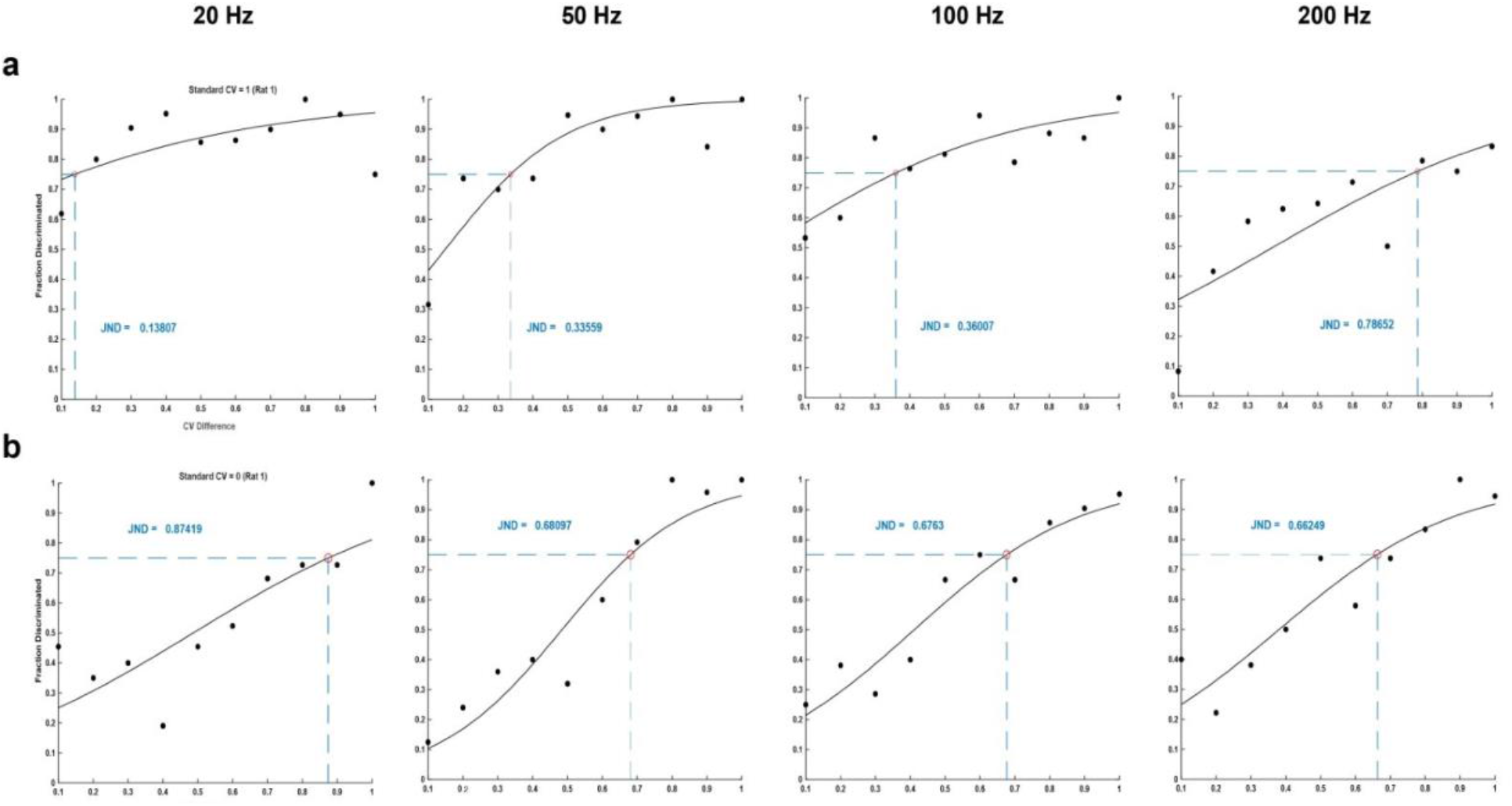
Examples of the sigmoidal curves used to find just noticeable differences (JNDs) for a single rat (rat 1) for the four frequencies tested. JND was taken as the CV difference value at the 75% correct mark on each sigmoidal curve. a) Fraction of trials correctly discriminated from a standard CV of 1. b) Fraction of trials correctly discriminated from a standard CV of 0. The circles in panels a and b indicate the fraction of correctly discriminated trials for each CV difference from the standard CV for that session.

**Figure 6.**
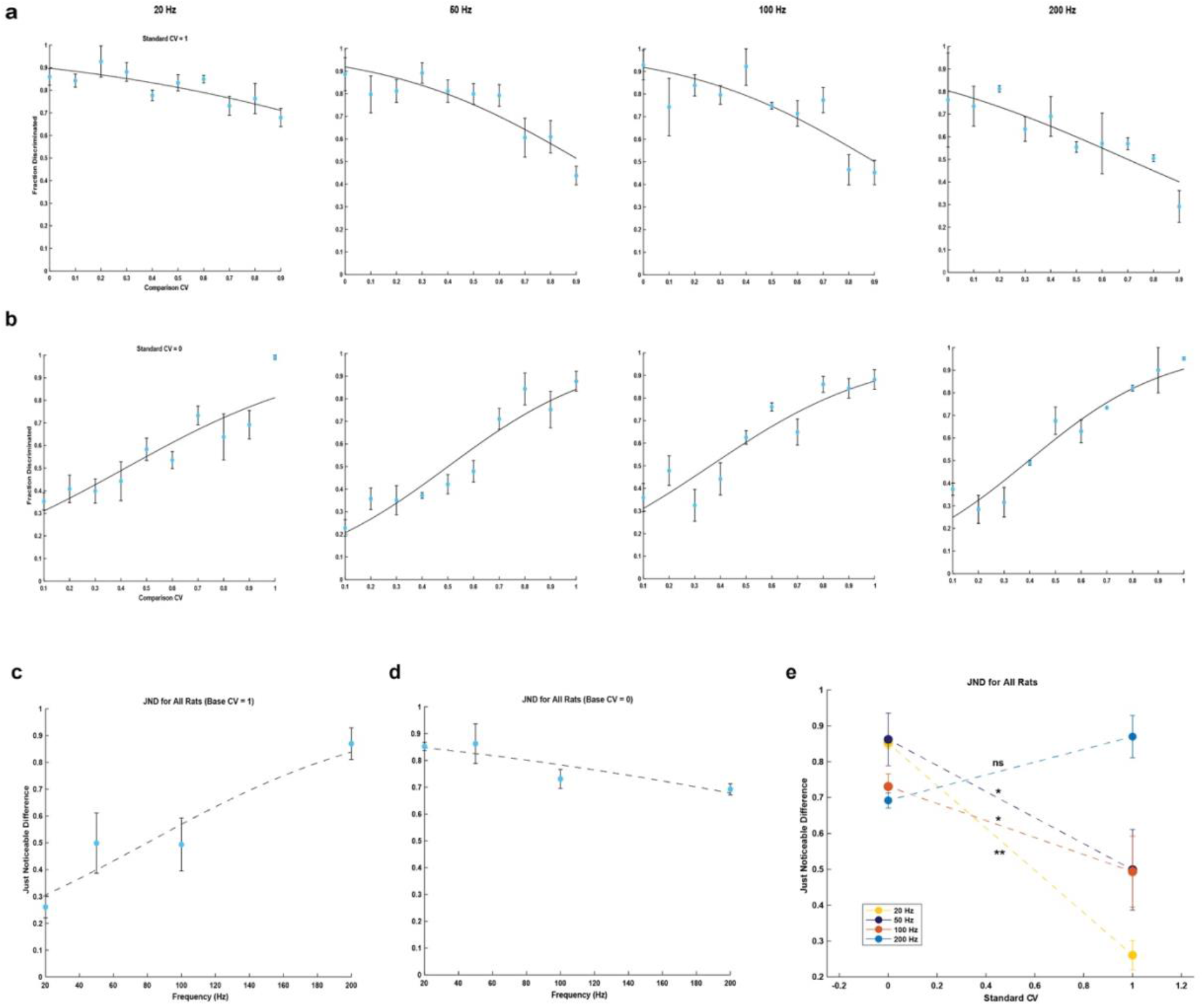
Average fractions of correctly discriminated trials and JNDs compared to higher (more aperiodic) and lower (more periodic) ranges of CV values across all rats. a) Fraction of correct trials discriminated at 20, 50, 100, and 200 Hz (n = 4, 4, 4, 2) compared to a lower CV value (more periodic stimulus). b) Fraction of correct trials discriminated at 20, 50, 100, and 200 Hz (n = 4, 4, 4, 2) compared to a higher CV value (more aperiodic stimulus). c, d) JND as a function of frequency. e) * and ** indicate a statistically significant (p<0.05 and p<0.01) difference between JNDs at standard CVs of 0 and 1 based on a paired t-test. ns indicates no statistical significance. a-e) Circles and error bars indicate mean ± s.e.m. Curves are sigmoid fits to the data.

Across all rats, the JNDs corresponding to the stimuli with a standard CV of 1 were significantly lower than those with a standard CV of 0 at all frequencies except 200 Hz (Fig. 6e). In addition, JNDs in CV increased as frequency increased for standard CV of 1 (Fig 6c). In contrast, JNDs in CV decreased as frequency increased for standard CV of 0 (Fig 6d). These results demonstrate the unique ability of rats to discriminate subtle differences in the temporal pattern of stimulation and establish that periodicity is an important parameter for generating distinct sensory perceptions.

## Discussion

Building on our previous work showing that SCS can encode artificial sensory perceptions (8, 9), in this study, we investigated the ability of irregularly patterned SCS to evoke sensory perceptions by delivering pulse trains that varied in the degree of aperiodicity. Rats learned to detect and discriminate aperiodic pulse trains, which suggests that temporally-varying SCS pulse trains can be meaningfully interpreted by the brain.

Sensory detection thresholds showed a significant decreasing trend with increasing aperiodicity at 20 and 50 Hz, but not at 10, 100, or 200 Hz. This suggests that the temporal pattern of SCS is a useful parameter for modulating perceptions within certain frequency ranges and might not be applicable at all frequencies. It is also possible that certain aperiodic temporal patterns generated a higher instantaneous frequency which could facilitate sensory perception at lower amplitudes. In a previous study in which frequency and duration were simultaneously varied at a constant amplitude, rhesus monkeys could detect stimulation trains with very small durations at higher frequencies. For instance, only 2-3 pulses were required to generate a sensory percept at 200 Hz and above frequencies. In the context of our study, template patterns for 1 CV at 20 Hz and 50 Hz had approximately 23% and 27% pulses occurring at intervals less than 5 ms (200 Hz), respectively (Fig. 7a). No pulses less than 5 ms occurred for 10 Hz while only 5% pulses <20 ms were observed at 0.8 CV (Fig. 7b). This may provide an explanation for the absence of decreasing trend in overall detection thresholds as CV was varied at 10 Hz but an unexpected deviation at 0.8 CV. Based on this evidence, a significant deviation from an average low-frequency stimulus due to aperiodic patterns could explain the decreasing amplitude trend at low frequencies but not at higher frequencies.

**Figure 7.**
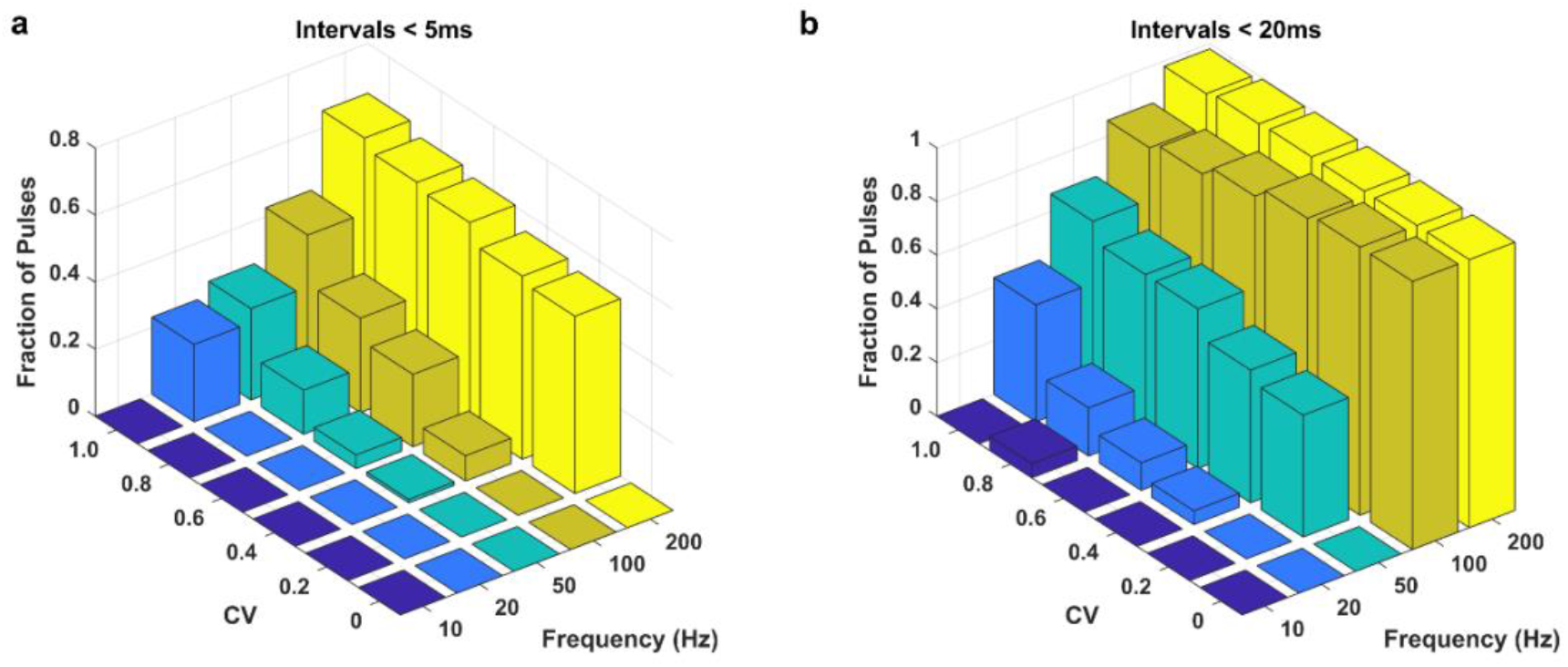
Pulse-count quantification of template patterns across CV and frequency for the two-second stimulation window. Z-axis displays the fraction of pulses that occurred at intervals less than a) 5ms (200Hz) and b) 20ms (50Hz).

All rats learned to discriminate between periodic (0 CV) and aperiodic stimuli (1 CV) at all frequencies. Two rats started discrimination at 50 Hz while two others started at 20 Hz (Fig 4, Supplementary Fig 1) before moving on to other frequencies. JNDs in CV from aperiodic stimuli (1 CV) were consistently lower than JNDs in CV from periodic stimuli (0 CV) for all rats at all frequencies except at 200 Hz. This indicates that the difference in randomness required to discern from highly aperiodic stimuli is smaller than that required from highly periodic stimuli. It appears counterintuitive because one would assume that any slight variation from periodic stimulation would be easily discernible. However, this result might highlight that the brain can easily decode variations from a highly random pattern compared to a more regularized pattern which might be perceived as non-naturalistic at lower frequencies. These differences in JNDs were reversed at frequency discrimination at 200 Hz (Fig 6c), suggesting that it becomes more difficult to discriminate between aperiodic patterns at higher frequencies.

At the beginning of each discrimination session for each rat, a threshold amplitude was determined at 0 CV and kept constant throughout the session when the CVs were randomized between 0.1-1 or 0-0.9. Based on the result that variation in stimulus aperiodicity impacts sensory detection thresholds (Fig 2), it is feasible that rats were discriminating the perceived difference in stimulation intensity instead of interpreting the temporal pattern. To challenge this explanation, stimulation must be delivered at the threshold amplitude of each CV value as it randomizes throughout the session. However, conducting that in the context of a rodent study may not be feasible and might need a parallel investigation in human subjects with SCS.

Future experiments could explore simultaneous modulation of frequency and periodicity. Previously, it was shown that, in rodents, SCS frequency discrimination followed Weber’s law at periodic stimulation i.e. when stimulation was delivered at 0 CV (9). It would be beneficial to determine whether JNDs in frequency follow Weber’s law at some or all CV values or if Weber’s law applies to JND in CV at certain or all frequencies. Modulating the frequency and periodicity of SCS simultaneously can greatly expand the number of distinguishable percepts available within a fixed frequency range. But, care must be taken to prevent patterns with high instantaneous frequency to avoid significantly altering the detection thresholds.

In conclusion, we successfully demonstrated that the temporal pattern of SCS is an important parameter that greatly impacts the detection and discrimination of sensory perceptions. The periodicity of a stimulation train should be further explored to generate reliable and distinguishable perceptions. Our current and previous results strongly indicate that SCS could effectively be used as a sensory neuroprosthetic technology to deliver artificial sensory feedback in clinical applications of BMIs.

## Acknowledgments

This research was supported by Indiana University Neurological Surgery research funding, Stark Neurosciences Research Institute research funding awarded to Amol P. Yadav, IUPUI University Fellowship awarded to Jacob C. Slack, and IUPUI UROP Scholarship awarded to Sidnee L. Zeiser.

## Author Contributions

Conceptualization: JCS, APY

Methodology: JCS, APY

Data collection: JCS, SLZ

Visualization: JCS, SLZ

Writing—original draft: JCS, SLZ, APY

Writing—review & editing: JCS, SLZ, APY

Supervision: APY

## Supplementary Information

**Supplementary Figure 1:**
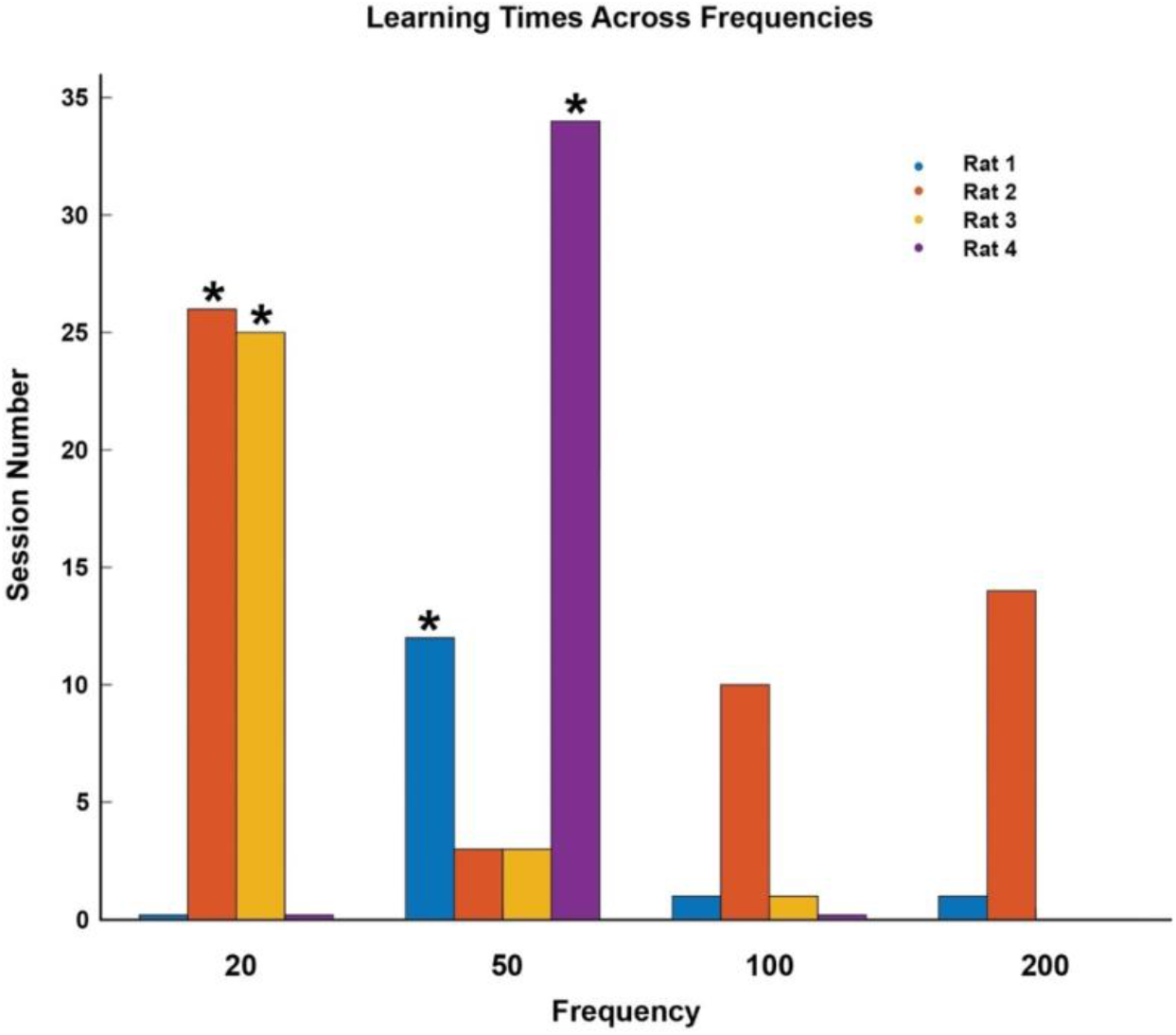
Discrimination learning times across frequencies. Time required to learn the initial discrimination task i.e., discrimination of periodic versus aperiodic stimulation patterns greatly varied across rats and frequencies. Rats 1 and 4 learned to discriminate periodicity firstly at 50 Hz while rats 2 and 3 learned it at 20 Hz before moving on to other frequencies. * indicates the frequency at which each rat began initial discrimination learning.

**Supplementary Table 1:** Electrode implant specifications for each rat. This study represents SCS implants that lasted 2-14 months in 6 rats. On average each rat performed ~253 trials per session for a total of ~43569 trials over ~9.2 months.

| Rat | Duration of Implant (months) | Average Trials per Session | Total Number of Trials |
| --- | --- | --- | --- |
| 1 | 14 | 297 | 71271 |
| 2 | 14 | 266 | 69606 |
| 3 | 12 | 243 | 61910 |
| 4 | 2 | 301 | 12624 |
| 5 | 6 | 142 | 16908 |
| 6 | 7 | 267 | 29092 |

